# Considerations for imaging thick, low contrast, and beam sensitive samples with liquid cell transmission electron microscopy

**DOI:** 10.1101/380923

**Authors:** Trevor H. Moser, Tolou Shokuhfar, James E. Evans

## Abstract

Transmission electron microscopy of whole cells is hindered by the inherently large thickness and low atomic contrast intrinsic of cellular material. Liquid cell transmission electron microscopy allows samples to remain in their native hydrated state and may permit visualizing cellular dynamics in-situ. However, imaging biological cells with this approach remains challenging and identifying an optimal imaging regime using empirical data would help foster new advancements in the field. Recent questions about the role of the electron beam inducing morphological changes or damaging cellular structure and function necessitates further investigation of electron beam-cell interactions, but is complicated by variability in imaging techniques used across various studies currently present in literature. The necessity for using low electron fluxes for imaging biological samples requires finding an imaging strategy which produces the strongest contrast and signal to noise ratio for the electron flux used. Here, we experimentally measure and evaluate signal to noise ratios and damage mechanisms between liquid and cryogenic samples for cells using multiple electron imaging modalities all on the same instrument and with equivalent beam parameters to standardize the comparison. We also discuss considerations for optimal electron microscopy imaging conditions for future studies on whole cells within liquid environments.

## 1 Introduction

Transmission electron microscopy (TEM) and scanning transmission electron microscopy (STEM) have become increasingly important tools for determining high resolution structures of biological samples. Cryogenic electron microscopy (cryo-EM) in particular has been a valuable technique for structural biologists due to its ability to resolve structures of individual proteins (*1*), viruses (*2*), and whole cells (*3*) at high resolutions. While time resolved cryo-EM and cryo electron tomography studies can be performed on semi-equivalent samples, these approaches cannot visualize dynamic processes from the same sample due to the static nature of freezing. Recent advances in instrumentation has allowed for electron microscopy of liquid samples (*4*), a technique that may permit visualizing physiological structures (*5, 6*) or dynamics (*7*) at high resolutions and in real-time. Liquid cell transmission electron microscopy (LC-TEM) has also been used to image whole cells in a fully hydrated state, where eukaryotes (*8–11*) and prokaryotes (*12–14*) have been imaged using both TEM and STEM. However, sample thickness is a consistent issue encountered when performing LC-TEM and is particularly problematic for large samples such as cells which are typically a minimum of 300-400 nm in thickness and can reach 1000 nm and greater depending on the cell type. Unfortunately, the penetrating power of electrons practically limits conventional TEM imaging to cells within this 300-1000 nm size range. While transmission of electrons and image formation is possible with these thickness ranges, imaging artifacts such as chromatic aberration in TEM, and beam broadening and narrow depth of focus in STEM can affect the final quality of the images formed (*15, 16*). Further complications arise from the inherent low contrast of biological samples due to their low atomic number and similar density to the media (water, buffer, or ice) that surrounds them. These factors result in images which have poor signal to noise ratio (SNR) even under ideal scenarios.

The electron irradiation sensitivity of biological samples additionally necessitates low electron fluxes for image formation during LC-TEM, likely at reduced values than those established for cryo-EM (*17*), which further decreases SNR and contrast. As such there is a need to optimize imaging conditions such that SNR, contrast, and resolution are maximized for images collected from thick samples with low electron fluxes. While this has been demonstrated experimentally for resin embedded cells or cryogenic thin sections (*16, 18*), and discussed in theory for liquid and cryogenic samples (*19–22*), at the time of this writing the optimal imaging modalities have not been demonstrated experimentally for LC-TEM across equivalent samples on the same instrument. Furthermore, despite demonstrations of hydrated cells imaged with LC-TEM in the literature, images collected with LC-TEM do not consistently show the same level of detail as those imaged with cryo-EM. As such, a comparison of cells imaged with LC-TEM and cryo-EM is needed to understand this discrepancy. In this manuscript we compare images of the organism *Cupriavidus metallidurans* acquired using multiple different imaging modalities and equivalent thicknesses for cryo-EM and LC-TEM to determine optimum imaging modalities for LC-TEM of biological samples.

## 2 Methods

### 2.1 Cell Culture

*Cupriavidus metallidurans* was obtained from ATCC and maintained in a standard nutrient broth (NB) media (3g beef extract and 5g peptone) supplemented with 50 μM AuCl at a pH of 6.8. Cultures were grown at 30°C with constant shaking and passaged 12 hours before EM experiments. Cultures were harvested by centrifuging 5mL of culture material at 3000g for 10 minutes and resuspending in 750 μL of NB media. Additionally, a culture was maintained in NB media without AuCl for comparison experiments.

### 2.2 Electron Microscopy

All electron microscopy was performed on a probe corrected 300kV FEI Titan equipped with a monochromator and GIF energy filter. The instrument was calibrated in TEM and STEM imaging modes for consistent electron fluxes for each imaging regime. Images collected for BF-TEM imaging for LC-TEM and cryo-EM was performed in TEM imaging mode with a 150 μm condenser aperture, a 40 μm objective aperture, with a beam current of 3.3 nA, a beam diameter of 11.45 μm, and a 1 second exposure for an image electron flux of 1 e^-^/Å^2^. The magnification for BF-TEM imaging was 4400 times magnified for a field of view of 4.45 μm. Images collected for BF-STEM and HAADF-STEM imaging were collected in STEM imaging mode with a beam current of 21 pA, a dwell time of 3.2 seconds, and a pixel size of 2.08 nm for an image electron flux of 1 e^-^/Å^2^. The magnification for STEM BF and HAADF imaging was 20,000 times magnified for a field of view of 4.27 μm. The semi-convergence angle for high convergence angle STEM imaging was 17.8 mrad, while the semi-convergence angle for low convergence angle STEM imaging was 5 mrad. EFTEM was performed with identical beam conditions to BF-TEM imaging for an electron flux of 1 e^-^/Å^2^, with a 30 eV energy slit centered on the zero-loss peak.

Cryo electron microscopy was performed with a Gatan 626 cryo holder. *C. metallidurans* cells were harvested after 12 hours of growth and concentrated via centrifugation. Three microliters of cells were frozen on Quantifoil EM grids with a Leica EM GP plunge freezer at 85% relative humidity, 25°C chamber temperature, −174°C ethane temperature, 30s preblot, and 2s blot time. Cells were imaged at an electron flux of 1 e^-^/Å^2^ for each imaging modality utilizing low dose imaging techniques.

Liquid cell electron microscopy was performed with a Hummingbird Liquid Stage. Cells were harvested after 12 hours of growth and concentrated via centrifugation, and 0.3 μL were deposited on a silicon device containing free standing silicon nitride membranes with a thickness of 10 nm. A second silicon device with another set of 10 nm membranes was placed on top of the first sandwiching the cells between them and hermetically sealed at the tip of the Hummingbird liquid stage. Cells were imaged at an electron flux of 1 e^-^/Å^2^ for each imaging modality utilizing low dose imaging techniques.

Electron energy loss spectroscopy measurements were taken for both cryo-EM and LC-TEM experiments near cell locations. Spectra were acquired with a 5 mm aperture and a dispersion of 0.25 eV with no energy shift so that the zero-loss peak was obtained with the plasmon in a single spectrum. Thickness in inelastic mean free paths was calculated using the log ratio technique for comparison of thicknesses between samples.

## 3 Results

While a number of different TEM imaging modalities exist, the vast majority of reports on biological structures to date have been performed with bright field TEM (BF-TEM), where contrast can be further improved with the use of phase plates (*23*) and energy filtered TEM (EFTEM). Although BF-TEM produces images with high contrast and possesses large depth of focus, for very thick samples it suffers from chromatic aberration which degrades image resolution (*15*). Low convergence angle bright field scanning transmission electron microscopy (BF-STEM) has been demonstrated to provide advantageous SNR and contrast over high angle annular dark field STEM (HAADF-STEM) and BF-TEM for thick resin embedded or cryo samples while maintaining good depth of focus (*16*). While classic HAADF-STEM imaging does not suffer from chromatic aberration, high convergence angles used typically for high resolution STEM imaging result in a narrow depth of field that can cause defocus artifacts in very thick samples, in addition to beam broadening artifacts that also degrade resolution for features near the bottom of a thick sample (*19, 20*). Experimentally determining which of these imaging modalities gives the best signal, contrast, and resolution for a given electron flux is needed to optimize results for whole cell imaging.

In order to accurately compare differences between imaging modalities, the role of changing sample thickness on resolution, contrast, and SNR must be considered. While patterned spacers are commonly used to dictate the minimum thickness of an assembled liquid cell, the tendency for inhomogeneous samples, such as cells, to clump together (or for environmental contaminants to be present on the surface of nanofluidic devices used to create the liquid cell) often results in a high degree of variability of the actual thickness of the liquid sample. **Supplemental Figure 1** illustrates this variability, where each image was acquired from separate samples each assembled with 300 nm spacers. Although both the original sample and nanofluidic devices were kept constant (and theoretically identical), BF-TEM images indicate the presence of increasing thickness between the three samples (**Supplemental Figure 1a**). Visually, features such as cell membranes and internal structures are lost as the thickness of the liquid layer increases. **Supplemental figure 1b** shows the corresponding low magnification BF-TEM images of the windows illustrating the loss of contrast and signal for increasing sample thickness even at low resolution. The discrepancy in image quality shown here necessitates that for comparison of various imaging modalities the liquid thickness for each image must be quantified to ensure that images being compared are from semi-equivalent thicknesses to improve interpretability. Therefore, for the data sets described below we collected electron energy loss spectrums (EELS) from locations near the cell positions immediately following the acquisition of high-magnification image sets. This permitted thickness to be quantified for every image area using the log ratio method where the thickness is expressed in terms of inelastic mean free paths (IMFP) (*24–26*). At the accelerating voltage used for these experiments, 1 IMFP in pure water with two 10 nm silicon nitride membranes corresponds to ~180 nm thickness. This approach allows for quantitative comparison between separate samples allowing for confidence that differences in SNR and contrast are not the result of changing thicknesses between samples.

### 3.1 Imaging Cells with Different Electron Imaging Modalities

Imaging was performed on *C. metallidurans* in cryo and liquid conditions for BF-TEM, EFTEM, high convergence angle STEM (α = 17.8 mrad), and low convergence angle STEM (α = 5 mrad). Representative images of cells at equivalent liquid/ice thicknesses are shown in **Figure 1**. The field of view changes slightly between different imaging regimes (due to magnification differences between STEM detectors and TEM CCDs) but all cells were imaged with an electron flux of 1e^-^/Å^2^ at the sample plane. **Figures 1a-d** are of cells imaged with LC-TEM, while **figures 1e-h** are of cells imaged with cryo-EM

**Figure. 1:**
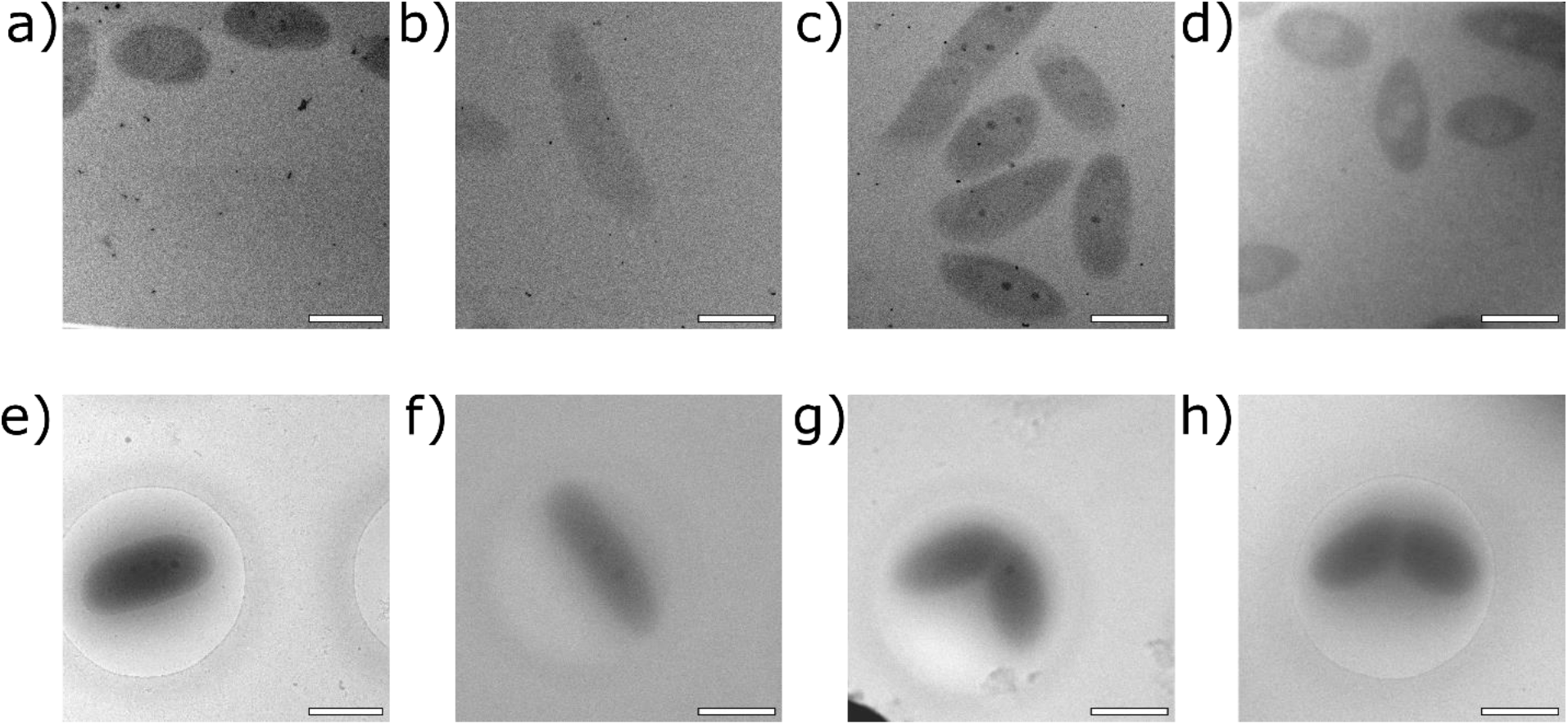
Comparison of whole cells of *Cupriavidus metallidurans* imaged with LC-TEM (a-d) and cryo-EM (e-h). (a, e) BF-TEM images with a 40 μm objective aperture. (b,f) High convergence angle (17.8 mrad) BF-STEM. (c, g) Low convergence angle (5 mrad) BF-STEM. (d, h) Energy filtered TEM with a 30 eV slit centered on zero loss peak. All scale bars are 1 μm.

**Figure 1a** shows cells imaged with BF LC-TEM where the thickness of the liquid measured near the cells was 1.98 IMFP. Some large structures of lower density and several smaller higher density structures associated with the cells are visible. **Figure 1b** shows cells imaged with high convergence angle BF-STEM in a liquid cell where the thickness was 2.93 IMFP and the convergence angle was 17.8 mrad which was typical for high resolution STEM imaging on the instrument. This was the thickest sample imaged and had the poorest contrast of all the images. The cell is only weakly visible against the background, although some small high density structures are visible. Prior to collection of this image, the cells were almost not visible at low magnifications when searching for cells to image, increasing the difficulty of accurately targeting a cell for imaging. **Figure 1c** shows a low convergence angle BF-STEM image of cells acquired in a liquid cell which had a measured thickness of 2.29 IMFP. The convergence angle was 5 mrad, which was the smallest angle that could be reached with the condenser aperture on the microscope. Cells visually appear to have stronger contrast than the cells acquired with high convergence angles, and both low and high density structures are visible with the cells. The cells were more easily detected during low magnification searching (as compared to identical scan parameters used for high convergence angle STEM) allowing for better targeting of imaging locations. **Figure 1d** shows an energy filtered image of cells in a liquid cell with a measured thickness of 2.14 IMFP. The energy slit used for this image was 30 eV in width and was centered on the zero-loss peak (ZLP). The cells visually have good contrast against the background and some low density structures are visible. The reference EELS spectra for each image are shown in **supplemental figure 2**. For each imaging strategy, imaging of cells was repeated but with cells frozen in vitrified ice.

In comparison to cells imaged with LC-TEM, **Figure 1e** shows a BF-TEM image of a cell frozen in ice with a thickness of 1.58 IMFP. This was the thinnest sample imaged, and the lipid bilayer membrane is visible surrounding the cell along with high density structures associated with the cells. **Figure 1f** shows a high convergence angle BF-STEM image of a frozen cell with a measured thickness of 2.03 IMFP. The image visually appears to show the least detail out of all the cryo-EM images although some high contrast structures are visible. Figure 1g shows a low convergence angle BF-STEM image of cells where the ice thickness near the cell was 2.27 IMFP. High density structures are visible associated with the cell although the cell edge is not as sharp as seen in the BF-TEM image. **Figure 1h** shows an energy filtered image at the same imaging conditions as figure 1d with a measured ice thickness of 1.92 IMFP near the cell. The EELS spectrums acquired for each sample and used for thickness calculation are shown in **supplemental figure 2**. While the cryo-EM BF-TEM was the thinnest and the LC-TEM high convergence angle BF-STEM image was the thickest, most images were obtained in nearly equivalent imaging regions around 2-2.5 IMFP. From comparing these cryo-EM images to their LC-TEM counterparts, it appears that the cryo-EM images generally have better contrast and SNR than the LC-TEM images despite similar sample thickness.

While apparent differences between electron imaging modalities can be seen by the images compared in **figure 1**, the magnitude of these differences as displayed visually can depend on the brightness and contrast settings that are chosen by the individual assembling the data. In order to avoid potentially subjective representation of differences between images, the SNR was quantified for each image to provide a measurable comparison of image quality. SNR was estimated using a single image autocorrelation technique which has been described previously in other fields (*27*). In short, the twodimensional autocorrelation function (ACF) was calculated for the image, where the maximum ACF value (at the origin) corresponded to the noise energy of the image. The noise free signal energy of the image was estimated by fitting a gaussian curve to the ACF values from the nearest neighbors to the ACF maximum around the origin. Once the noise energy, signal energy, and the mean value of the image has been determined the SNR can be estimated (*27*). To be sure of the accuracy of the method for estimating SNR a standard STEM image was duplicated and white gaussian noise with a known increasing variance was added to each image. SNR values were then estimated for each of the simulated images, where it was observed that the estimated SNR decreased as noise was added (**supplemental figure 3**). This confirms that the single image autocorrelation technique for SNR determination properly estimates the image SNR and our implementation of the previously published methods worked as described.

**Figure 2a** shows a cropped version of figure 1a used for SNR estimation from a BF-TEM image taken with LC-TEM. Images were cropped around just the cell as, especially for cryo-EM samples, the ice thickness is often thinner in areas away from the cell. As EELS measurements were taken as near to the cells as possible and changes in thickness can have significant impact on SNR values only image regions which were likely to be near the measured thickness were used for SNR estimation. **Figure 2b** shows the two-dimensional ACF calculated for **figure 2a**, and **figure 2c** shows a plot of ACF values 4 pixels on either side of the maximum of the ACF across the x-axis for ease of visualization. The orange line in figure 2c is the ACF values where the maximum value is the noise energy, while the blue line is the gaussian curve fit to the data where the value of the gaussian fit at the origin is used as the signal energy for SNR estimation. **Figure 2d-f** shows the same as **a-c** but for BF-TEM images obtained with cryo-EM. **Table 1** shows the SNR values for each image corresponding to the images in **figure 1**. Cryo-EM images had higher SNR values than their corresponding LC-TEM counterparts, and BF-TEM and EFTEM had the highest measured SNR of all the imaging modalities. HAADF images were also simultaneously acquired for both high and low convergence angle data sets and SNR values were estimated for HAADF images from the same cropped regions as for their BF counterparts. For both low and high convergence angle HAADF-STEM the estimated SNR was higher in the BF images. Images collected for HAADF-STEM are shown in **supplemental figure 4**.

**Figure. 2:**
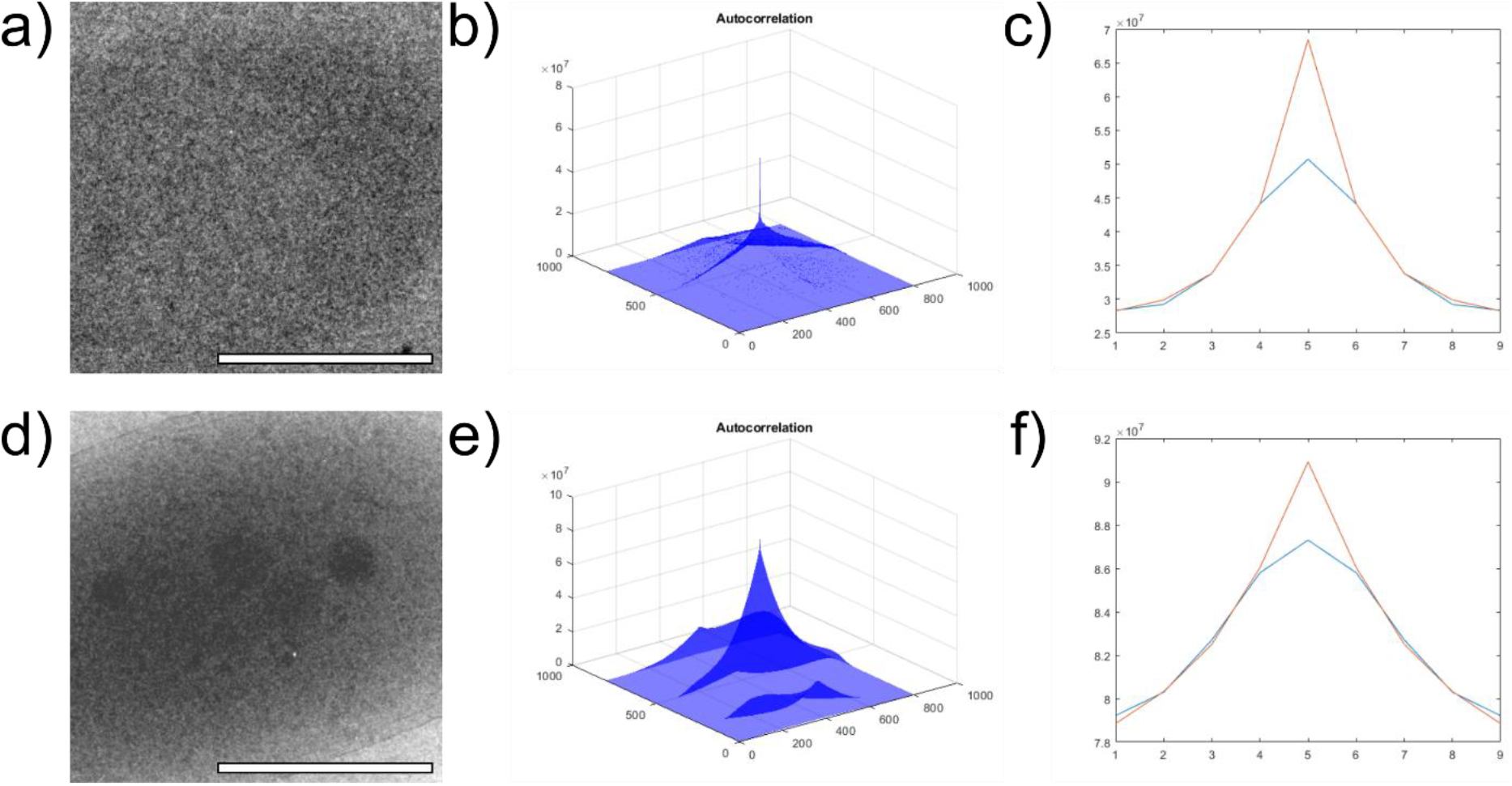
a) 400×400 pixel crop of **figure 1a** used in SNR estimation. b) Two-dimensional ACF calculated from (a). c) Sample of ACF curve along the x-axis (orange) and the gaussian curve (blue) used to estimate the signal energy of the image. d) 400×400 pixel crop of figure 1e used in SNR estimation. e) Two-dimensional ACF calculated from (d). f) Sample of ACF curve along the x-axis (orange) and the gaussian curve (blue) used to estimate the signal energy of the image. All scale bars are 500 nm.

**Table 1:** Estimated SNR Values from LC-TEM and Cryo-EM Images

| | BF-TEM | BF-STEM<br>$\alpha = 17.8$ mrad | HAADF-STEM<br>$\alpha = 17.8$ mrad | BF-STEM<br>$\alpha = 5$ mrad | HAADF-STEM<br>$\alpha = 5$ mrad | EFTEM |
| --- | --- | --- | --- | --- | --- | --- |
| LC-TEM | 1.8116 | 0.4892 | 0.4533 | 0.5683 | 0.5229 | 12.0492 |
| Cryo-EM | 16.7694 | 1.7283 | 0.8921 | 3.9663 | 1.1996 | 15.9064 |

$\alpha$ = semi-convergence angle of the STEM probe

### 3.2 Comparison of Damage Between Cryo-EM and Liquid-EM

The role of electron beam damage has been well characterized for cryo-EM, where a relationship between cumulative electron flux and loss of structure has been described in detail (*28–31*). Despite this knowledge it remains unknown how a loss of structure impacts the functionality of biomolecules, whether the catalytic mechanism of an enzyme, the protein coding function of an mRNA, or the transport, recognition, and binding of a transcription factor. Dehydrated catalase proteins have been demonstrated to lose their catalytic function at electron flux rates as low as 0.05 e^-^/Å^2^ (*32*). However, it remains unclear how similar a hydrated protein sample may behave in LC-TEM due to the added secondary damage mechanisms from radiolysis. For tomography of whole cells with cryo-EM, optimal cumulative electron fluxes are generally around 100 e^-^/Å^2^, but can be up to as much as 200 e^-^/Å^2^, and achieve resolutions of 20-60 Å (*28*).

However, cumulative electron fluxes in cryo-EM above 100 e^-^/Å^2^ often begin to cause bubble formation from hydrogen bubble generation which can cause visible image distortions (*28*). We have previously reported beam driven shrinking of *C. metallidurans* in LC-TEM for cumulative electron fluxes considerably lower than those described for cryo-EM of whole cells at 1-4 e^-^/Å^2^ (*17*). These cells had been cultured in media which had been supplemented with 50 μM AuCl. As others have postulated that metals present in liquid or frozen media could act as scattering centers that might locally increase electron beam induced damage (*33, 34*) we considered that the presence of the AuCl in the media may have caused the lower beam tolerance observed previously. We therefore repeated LC-TEM imaging with and without the presence of gold chloride in the media to determine if the presence of low concentrations of heavy metals in solution can noticeably accelerate secondary damage. We also repeated these damage series experiments with cryo-EM, both in the presence of gold chloride and without, to compare the onset of electron beam driven damage and the types of morphological changes that are observed between LC-TEM and cryo-EM. **Figure 3** shows BF-TEM damage series of *C. metallidurans* in increments of 1 e^-^/Å^2^ for a total cumulative dose of 4 e^-^/Å^2^ by the fourth image in the series. **Figures 3a** and **3b** are of cells in a liquid environment, where **3a** are cells grown with supplemented gold and **3b** are cells which have not been grown in gold supplemented media. The final column in **figure 3** shows the first and last image from the damage series aligned and overlaid on top of one another allowing for comparison of changes in the cells between images. Both LC-TEM damage series illustrate the same beam driven cell shrinking at equivalent onsets and similar to what has been reported previously (*12, 17*), indicating the presence of gold in the growth media does not considerably increase the beam driven morphological changes observed in the data. **Figures 3c** and **3d** show comparative damage series of *C. metallidurans* grown with (**3c**) and without (**3d**) gold, but imaged with cryo-EM rather than LC-TEM. During identical electron flux increments as used for LC-TEM above, visible morphological changes were not observed in the cells imaged with cryo-EM. However, similar cryo-EM damage series of *C. metallidurans* at much higher cumulative electron flux values showed the onset of visible bubble formation occurring around 300-400 e^-^/Å^2^, shown in **supplemental figure 5**, in agreement with other published results on damage observations in whole cells (*28*).

**Figure. 3:**
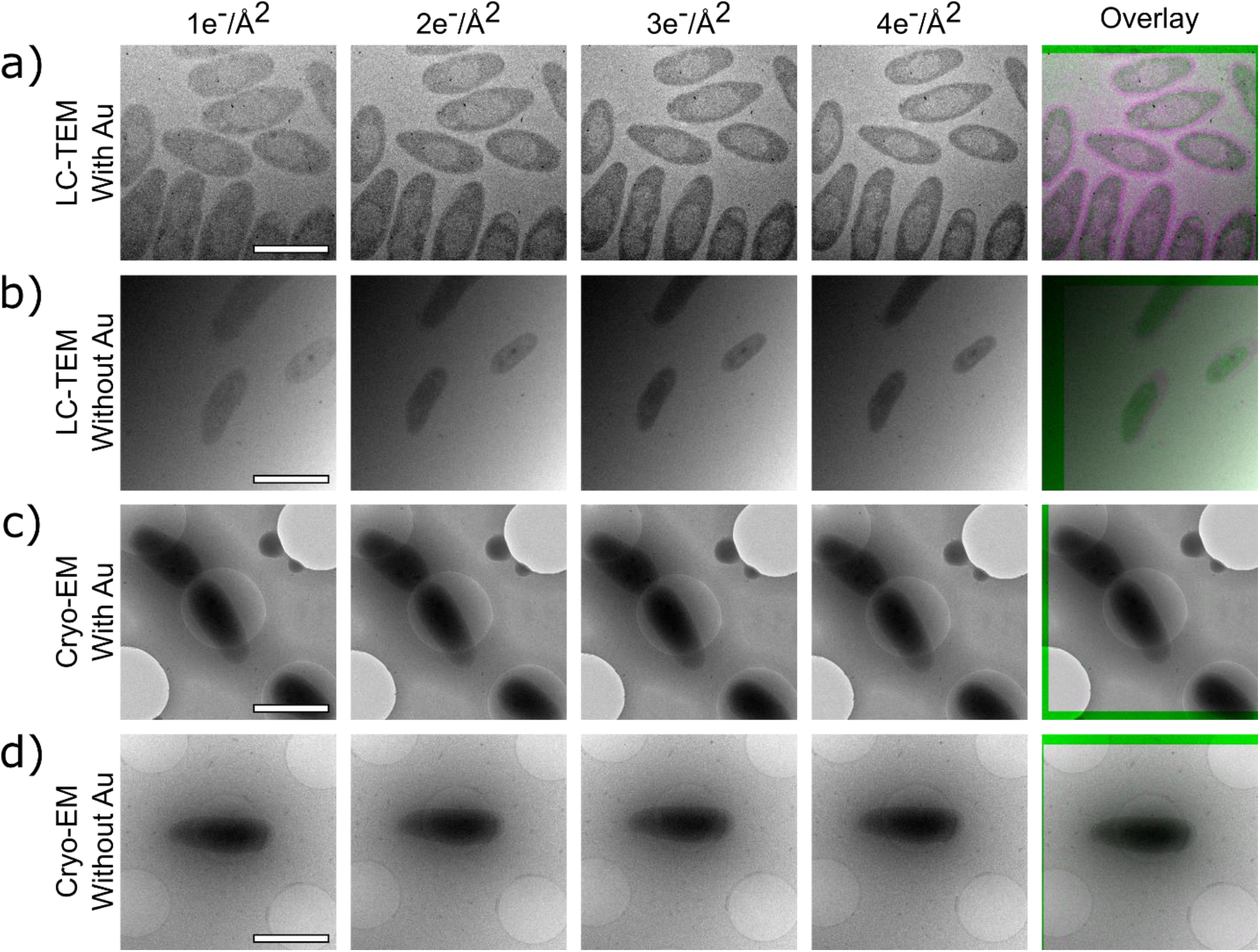
Comparison of electron irradiation of C. metallidurans cultured with or without the presence of AuCl between cryo-EM and LC-TEM. Each image was acquired with BF-TEM with a 40 μm objective aperture with a single image electron flux of 1 e^-^/Å^2^, and the time between each image was 20 minutes. The cumulative electron flux for the series is shown above each image in the series. The final column is an aligned and overlaid image of the 1^st^ and 4^th^ images to show changes in cell morphologies throughout the imaging series. Perfect gray scale overlay indicates now changes, whereas the presence of purple indicates an area of density that was present in the initial image, but that contracted by the final image. a) LC-TEM images of cells cultured in NB media supplemented with AuCl. b) LC-TEM image series of cells cultured in NB media. c) Cryo-EM image series of cells cultured in NB media supplemented with AuCl. d) Cryo-EM image series of cells cultured in NB media. All scale bars are 1 μm.

## 4 Discussion

For the comparisons made here it was important that thickness measurements were as close to equivalent as possible to ensure that observed differences between images were not an artifact of changes in sample thickness. It should be noted that the thicknesses presented here are not ideal for achieving maximum resolution due to limitations of LC-TEM sample preparation. While automated plunge freezing of liquid samples for cryo-EM has become increasingly reliable and controlled, reliable reproduction of samples of a desired thickness in LC-TEM remains challenging, especially with non-homogenous samples that have a tendency to clump together. The outward bulging of membranes under the pressure differential of the column further exacerbates this issue, where the thickness can increase substantially towards the center of the imaging area. Because of these imaging challenges with LC-TEM, it was found that the thicknesses for liquid samples were commonly thicker than ideal cryo-EM samples. Because of this, ice thicknesses for cryo-EM samples were intentionally made thicker than ideal for appropriate comparison to the LC-TEM data. We have previously shown LC-TEM results on *C. metallidurans* that show increased detail when compared with the data shown here when a very thin sample was achieved (*17*), although replication of these sample thicknesses across the multiple samples imaged here was not accomplished within the experimental time frame despite numerous attempts. As a result the data here is not representative of what may be maximally achieved for resolution and SNR with LC-TEM of biological samples, but a comparison of which imaging modalities give the strongest SNR for samples at comparable thicknesses. With improved instrumentation and sample preparation strategy it may eventually prove possible to overcome the current challenges with thickness reproducibility for LC-TEM.

Of the imaging modalities we tested, it was found that EFTEM and BF-TEM resulted in the highest SNR values compared to BF-STEM imaging at the thicknesses measured (near 2 IMFP). EFTEM with the energy slit on the ZLP appears to be slightly advantageous to BF-TEM, likely as the inelastically scattered electrons which do not contribute to image formation are removed from the image by the filter, improving the SNR. It is also possible to place the energy slit on the carbon edge or the most probable energy loss to collect signal scattered by only the cells in the image. The reduction in signal with this method however requires large integration times which considerably increases the electron flux on the sample (*15*). As LC-TEM of biological samples will likely be limited by the radiation tolerance of the cells imaged, increasing the incident electron flux used for image generation is not a practical strategy when considering options for image formation. While these methods provide the strongest SNR, we note that we have not quantified resolution between imaging modalities, where it would be expected that both BF-TEM and EFTEM imaging will show chromatic aberration effects at higher resolutions (*15*). For experiments requiring higher resolutions, simulations have shown that beam broadening resolution artifacts in BF-STEM are less significant than chromatic aberration artifacts in BF-TEM, suggesting STEM imaging may be preferable for imaging thick samples if resolution is preferred to SNR (*20*).

In addition to advantages of resolution, STEM imaging may be desirable for nanoparticle labeling of biomolecules of interest, as tracking of the positions or locations of biomolecules may require labeling with high contrast nanoparticles or quantum dots. Several groups have been exploring the possibility of using nanoparticle labeling with LC-TEM (*8, 10, 35, 36*) and typically use STEM imaging for this application. Interestingly, the BF-STEM images from high convergence angle STEM show the worst SNR for all the imaging methods tested, in addition to visually appearing to have the weakest contrast and show the least amount of detail. Contrast of cells at low magnifications was also considerably worse for high convergence angle STEM. This is important, as low dose imaging strategies to minimize electron irradiation of the sample prior to image collection commonly use low magnifications to target cells for imaging. Since cells were not often visible during low magnification searching with high convergence angle STEM, the final images were often acquired “blind,” where an image was acquired without knowing if a cell would be within the frame of view. On the other hand, cells imaged with low convergence angle BF-STEM were much more visible at low magnifications, such that regions of interest containing a cell or cells could be centered for accurate final image capture. It was also found that cells acquired with low convergence angle BF-STEM were more commonly in focus than images acquired with high convergence angles. While patterned grid bars were used for focusing prior to image acquisition (*17*), the narrow depth of focus of high convergence angle STEM has increased sensitivity to variations in sample position due to membrane bulging or small sample tilts laterally from where focus was captured. Therefore, if STEM imaging is preferred over TEM imaging our results show that low convergence angles should be used due to increased SNR, contrast, and depth of focus. It should be noted that the low convergence angle experiments here used a STEM probe with a 5 mrad convergence angle. While the instrument used for these experiments did not have it, a smaller condenser aperture can be used to obtain 1 mrad probes which will likely further improve SNR, contrast, and depth of focus.

Notably, the quantification of SNR values for each image suggests that despite potentially subjective visual interpretation of the images, LC-TEM images have consistently lower SNR values than their cryo-EM counterparts despite achieving similar thickness samples. One explanation for this discrepancy may be the presense of the silicon nitride membranes for LC-TEM samples. The cells imaged with cryo-EM were always over holes in a holey carbon film, while cells in LC-TEM always had two 10 nm silicon nitride membranes above and below the cell. It is possible the presence of these membranes, although thin, may cause increased noise signal in LC-TEM images. Previous HAADF-STEM image simulations of Pb nanoparticles has shown that the presence of a silicon nitride membrane can cause a decrease in SNR which scales with increasing membrane thickness (*37*). It may be that the presence of the membranes alone is enough to account for the decrease in SNR for otherwise similar samples. Alternatively, there may potentially exist physical differences in the structure of liquid versus frozen water (i.e. tunneling channels in vitreous ice) that contribute to the difference in SNR between cryo-EM and LC-TEM - although physical differences between ice and liquid interactions with electrons requires additional characterization.

Finally,_the LC-TEM damage experiments performed on *C. metallidurans* show what appear to be beam driven morphological changes regardless of the content of the liquid media the cells are grown in. This is similar to what has been shown previously both for *C. metallidurans* (*17*) as well as magnetotactic bacteria imaged with STEM imaging in liquid (*12*). When compared with cryo-EM, the beam driven sample changes are not observed until significantly higher cumulative electron fluxes. It is important to note that this difference does not necessarily indicate that damage is not occurring in the cryo-EM data sets, only that it is not manifested visibly at the resolutions achieved in the images. The frozen state of the sample in cryo-EM may result in damage to proteins and other cell components to be “locked” in place, whereas damage in liquid environments may be free to propagate through the biomolecule and the cell causing conformational changes at the molecular and cellular level. While loss of structural information with increasing electron flux has been well characterized for cryo-EM (*30*), it is not yet clear how a loss of structure corresponds to a loss or alteration of function for a protein or other biomolecule. As interpretation of physiological results from LC-TEM experiments on biological samples will depend highly on the ability to account for damage events, identifying the electron inactivation thresholds for biomolecules is a critical step towards accurate characterization of organisms and physiological processes with LC-TEM. Future work will focus on identifying these inactivation thresholds to determine what electron fluxes are required for imaging dynamics without significantly altering the natural state of the cell. With the new advances reported here and in recent publications by other groups, a cadre of tools are becoming available that position the field to start rigorously probing the beam effects on biomolecule structure and function to fully explore the potential for LC-TEM to visualize physiological dynamics in-situ.

## Acknowledgments

This work was supported by the Department of Energy’s Office of Biological and Environmental Research Molecules to Mesoscale Bioimaging Project #66382 and was performed using EMSL, a national scientific user facility sponsored by the Department of Energy’s Office of Biological and Environmental Research and located at PNNL. Effort for T.S. was supported by the National Science Foundation CAREER award (Grant No DMR-1564950).

## Author Contributions

J.E.E. conceived the work and J.E.E. and T.M. planned the experiments. T.M. performed all LC-TEM experiments. J.E.E. and T.M. analyzed the data and wrote the initial manuscript. T.S. assisted with data analysis and interpretation. All authors edited and approved the manuscript.

